# Expanding the list of sequence-agnostic enzymes for chromatin conformation capture assays with S1 nuclease

**DOI:** 10.1101/2023.06.15.545138

**Authors:** Maria Gridina, Andrey Popov, Artem Shadskiy, Nikita Torgunakov, Andrey Kechin, Evgeny Khrapov, Oxana Ryzhkova, Maxim Filipenko, Veniamin Fishman

## Abstract

This study presents a novel approach for mapping global chromatin interactions using S1 nuclease, a sequence-agnostic enzyme. We develop and outline a protocol that leverages S1 nuclease’s ability to effectively introduce breaks into both open and closed chromatin regions, allowing for comprehensive profiling of chromatin properties. Our S1 Hi-C method enables the preparation of high-quality Hi-C libraries, marking a significant advancement over previously established DNase I Hi-C protocols. Moreover, S1 nuclease’s capability to fragment chromatin to mono-nucleosomes suggests the potential for mapping the three-dimensional organization of the genome at high resolution. This methodology holds promise for an improved understanding of chromatin state-dependent activities and may facilitate the development of new genomic methods.

## Introduction

Genomic assays, which profile chromatin properties on a whole-genome scale, rely significantly on enzymatic activities corresponding to different chromatin states. For instance, MNA-seq assay utilizes micrococcal nuclease’s preferential digestion of nucleosome-free regions, while DNase hypersensitivity assay exploits DNase I enzyme’s preferential cutting in open chromatin regions [1]. The discovery of transposase Tn5 and similar enzymes, which also exhibit affinity to open chromatin regions, has further simplified the analysis of open chromatin [2]. Therefore, expanding the repertoire of enzymes with specific, chromatin-state-dependent activities is crucial for developing novel genomic methods.

In certain applications, enzymes with activity limited to specific chromatin states are necessary, while in others, uniform genomic coverage is essential. Chromatin conformation capture assays, including its whole-genome modification known as Hi-C, notably benefit from using enzymes with sequence-agnostic chromatin cleavage. Hi-C has been utilized by others and us to study genome architecture across various tissues [3] and individual cells [4], understand mechanisms underlying gene regulation in development [5,6], explore alterations in gene expression in cancer [7], assemble genomes [8,9], and identify structural variants [10].

The Hi-C technique involves several mandatory steps: cell fixation with formaldehyde to preserve native genome folding in the nucleus, chromatin fragmentation, chromatin end labeling by biotin, and chromatin ligation [11]. Consequently, fragments in close spatial proximity are ligated, and a biotin label is employed to selectively enrich these ligated fragments.

Enzymes for chromatin fragmentation in a Hi-C experiment can vary depending on the study’s purpose. Initial protocols recommended using restriction enzymes with a 6 bp recognition site, limiting data resolution to approximately 10 kb [12]. However, the introduction of 4-base cutters and other modifications significantly improved this resolution [13]. Genome-wide distribution of restriction sites and sequencing depth are the limiting factors for resolving fine-scale chromatin organization. These limitations can be circumvented by using a mixture of restriction enzymes or sequence-agnostic nucleases for chromatin fragmentation [14–16]. Currently, two sequence-agnostic enzymes, MNase and DNase I, are used in Hi-C studies. MNase Hi-C can generate nucleosome-level interaction maps and is proficient in loop detection, although it is less apt for compartment detection than conventional Hi-C [17]. Nevertheless, employing MNase for Hi-C is labor and time-consuming as it requires additional steps for cell fixation, meticulous optimization of MNase amount for uniform genome digestion, isolation of the dinucleosome fraction, and is unsuitable for analyzing chromatin devoid of nucleosomes. Using DNase I Hi-C protocol improves data resolution and genomic coverage over conventional Hi-C [15]. However, DNase I Hi-C libraries contain a high level of non-informative fragments or “dangling ends” (DE) [11], necessitating deeper sequencing.

S1 nuclease, a sequence-agnostic enzyme, degrades single-stranded nucleic acids and cleaves nick, gap, mismatch or loop structures in double-stranded DNA. S1 nuclease can also introduce breaks into double-stranded DNA at high enzyme concentrations [18]. S1 nuclease’s ability to cleave unpaired DNA is exploited in S1-seq and S1-END-seq for studying DNA secondary structure and meiotic double-strand break end resection on a genome-wide scale [19–21].

In this study, we delineate a profile of chromatin digestion by S1 nuclease and establish a protocol utilizing this enzyme for mapping global chromatin interactions. Our findings reveal that S1 nuclease effectively introduces breaks into both open and closed chromatin. The developed S1 Hi-C method enables the preparation of high-quality Hi-C libraries, surpassing the performance of the previously published DNase I Hi-C protocol. Furthermore, S1 nuclease fragments chromatin to mono-nucleosomes, implying that the three-dimensional organization of the genome can be mapped with high resolution.

## Methods

### Experimental procedures

#### Human cells isolation and culture

Human K562 cells were grown in RPMI-1640 medium with 10% FBS and a penicillin/streptomycin mix Human peripheral blood mononuclear cells (PBMC) were isolated from peripheral blood samples, which were collected in tubes with EDTA. RBC lysis buffer (BioLegend) was used for erythrocyte removal according to the manufacturer’s instructions. Isolated PBMCs were processed immediately for crosslinking.

#### S1 Hi-C protocols

1. Cell crosslinking described previously [11]
2. Cell lysis and chromatin fragmentation
  2.1. The pellet of cross-linked cells was placed on ice and gently resuspended in 1 ml of cell lysis buffer (10 mM Tris–HCl pH 8.0, 10 mM NaCl, 0.5% Igepal).
  2.2. The cells were incubated 1 hour with intermittent rotation at RT.
  2.3. Centrifugation was performed at 2500g for 5 min.
  2.4. The supernatant was removed, and the pellet was gently resuspended in 200 μl DNase buffer (50 mM Tris–HCl pH 7.5, 0.5 mM CaCl2) and 0.2% SDS.
  2.5. The cells were incubated at 37 °C for 1 hour.
  2.6. Control point #1: 5 μl lysed cells were saved to check gDNA integrity.
  2.7. SDS was quenched by adding 25 μl 10% Triton X-100 10 min at 37 °C.
  2.8. Centrifugation was performed at 2500g for 5 min.
  2.9. Pellet was resuspended in 500 μl 1x S1 nuclease buffer with 1% Triton X-100.
  2.10. Centrifugation was performed at 2500g for 5 min.
  2.11. Pellet was resuspended in 80 μl 1x S1 buffer
  2.12. S1 nuclease (200 U; Thermo Scientific) was added and incubated at 37 °C for 1 hour.
  2.13. The reaction was stopped by adding 5 μl 500 mM EDTA.
  2.14. Control point #2: 5 μl of the reaction was saved to check the S1 nuclease digestion efficiency. 90 microlitres of lysis buffer (10 mM Tris–HCl pH 8.0, 10 mM NaCl, 0.3% SDS) and 5 μl proteinase K (800 units/ml) were added to both controls. The controls were reverse cross-linked at 65 °C for at least 8 h. DNA was extracted by the standard phenol–chloroform method.
  2.15. Reaction was purified by 0.8 volume of AMPure beads according to the manufacturer’s recommendations.
  2.16. The beads were resuspended in 100 μl 1x NEBuffer 3.1, and the AMPure Beads remained in the mixture.
3. Biotin labelling, in situ ligation, cross-link reversal, removal of biotin from unligated ends and preparing libraries for NGS described previously [11].

We sequenced the S1 Hi-C libraries using paired-end reads with a length of 150 bp. Read depth was 50-100 k reads per sample for shallow sequencing, ∽37 mln reads for deeper sequenced S1 Hi-C on human K562 cells, ∽25 mln reads for S1 digested PBMC chromatin and ∽65 mln reads for K562 and digested chromatin samples.

### Computational data analysis

#### Public datasets

The list of public datasets analyzed in this study is provided in Supplementary Table 1

#### Hi-C data analysis

Hi-C data analysis and statistics computation was performed as described previously [11] with a slightly modified version of the Juicer 1.6 script and an altered computation of statistics. The script is publicly accessible on GitHub (https://github.com/genomech/juicer1.6_compact).

#### Chromatin cut sites analysis

Sequencing data obtained in this study or downloaded from public datasets were aligned to the hg38 human genome assembly using BWA v.0.7.17-r1188. Then, bam file sorting and indexing was performed using samtools. Read-pair fragment size distributions were calculated using bamPEFragmentSize from deeptools. The distributions were plotted using matplotlib.

BigWig files containing cut site coverage were generated using bamCoverage from deeptools with the options --OffSet 1 --binSize 1.

We processed obtained bigwigs and public data describing genomic features locations in bed-format (Supplementary Table 1) using a python script that enables plotting of the average cleavage signal within a window of -3000 to +3000 from the middle of the cutting sites of nuclease peaks, with (step size of 2 bp,) (refer to Figure 2B). The script is publicly accessible on GitHub (https://github.com/genomech/PlotBigwigOnBed). To ensure accuracy, we excluded genomic regions that are blacklisted by Encode for hg38 from the analysis (Supplementary Table 1). We similarly analyzed the mean S1 cleavage signal and other genomic features in the window of differentially expressed genes’ transcription start sites (TSS) using K562 RNA-seq data publicly available on Encode (Supplementary Table 1). To achieve this, we defined 5% of the most abundant transcripts based on transcript per million (TPM) as highly expressed genes, and a tier of 5-20 percentile of the most abundant transcripts as low expressed genes (i.e. top 20% excluding the highly expressed genes). Additionally, we selected 20% of transcripts with the lowest expression levels as the non-expressed ones. Finally, we used the same script to generate plots (refer to Figure 2B).

To identify S1 sequence specificity of the S1 cut sites, cut positions were obtained from the alignment files using the following script: “*samtools view -F 16 alignment*.*bam* | *awk ‘$6 ∽ “^[0-9]+M” && $0 !∽ “MD:Z:0[ACTG]” {print $3, $4}’*” for forward reads and “*samtools view -f 16 alignment*.*bam* | *awk ‘$6 ∽ “M$” && $6 ∽ “^[0-9]+M” && $0 !∽ “[ACTG]0[[:space:]]” && $0 !∽ “[ACTG]0$” {sum = $4 + 99; print $3, sum}’*” for reverse reads. Genomic regions flanking cut sites were extracted using pysam. Consensus logo for the extracted sequences was created using WebLogo 3.5.0.

#### Genome-wide read coverage depth analysis

Read coverage depth of Hi-C libraries prepared with S1, DNase I and DpnII was analyzed with bigWig files generated using bamCoverage from deepTools 3.5.1 with the option -- normalizeUsing RPKM [22]. As an example of a library prepared with DpnII we used publicly available GM12878 Hi-C data from 4D Nucleome Data Portal [23] (Supplementary Table 1). To build distributions of read coverage depth each individual chromosome in bigWig files was divided into segments of 500 nucleotides, from which 100,000 segments equidistant from each other were selected, followed by calculation of the coverage sums in each segment. Analysis was performed using pyBigWig 0.3.18 [22], NumPy 1.21.6 [23], the histograms of distributions were plotted using matplotlib 3.5.3 [24].

## Results

### Boosting Data Yield in Hi-C Protocol through DNase I Substitution with S1 Nuclease

Previously, we established a robust and efficient DNase I Hi-C protocol, yielding high-quality Hi-C maps [11]. Nevertheless, this method generated more dangling ends (DEs) compared to the traditional Hi-C approach. DEs represent non-chimeric DNA fragments that do not contribute valuable information for Hi-C analysis. These fragments are reduced during Hi-C library preparation through a following process: DNA ends are labeled with biotin after chromatin digestion; biotinylated nucleotide internalization following DNA end ligation; biotinylated nucleotides remaining on the unligated DNA ends are removed using exonuclease; molecules containing internal biotin (i.e. ligation products) are enriched by streptavidin pulldown. We suspect that the surplus of DEs in the DNase I Hi-C protocol is attributed to the nickase activity of DNase I. Following DNase I digestion, the 5’-3’ exonuclease activity of the Klenow Fragment elongates the nicks, incorporating biotin-dCTPs into the DNA molecule. Consequently, not only are the ligation products internally labelled with biotin, but also DNA molecules that have not participated in the ligation process

Like DNase I, S1 nuclease is a sequence-agnostic nuclease capable of cleaving dsDNA, nicks, and ssDNA. We theorized that these activities of S1 nuclease could prevent the generation of nicked DNA during chromatin digestion. To verify this, we first ensured that the S1 enzyme could digest PFA-fixed chromatin, producing a discernible digestion pattern (Fig 1A). Subsequently, we modified the cell lysis and chromatin fragmentation steps (see Methods) to make the Hi-C protocol compatible with the S1 chromatin digestion. Using this modified method, we prepared S1 Hi-C libraries for 16 human peripheral blood samples and the K562 human immortalized cell line. During libraries preparation, we examined the products of digestion and ligation steps and found that they satisfy Hi-C quality standards (Fig. 1A), although we note that the pattern of chromatin fragmentation by S1 nuclease looks slightly different for different cell types (Fig. 1A).

**Figure 1.**
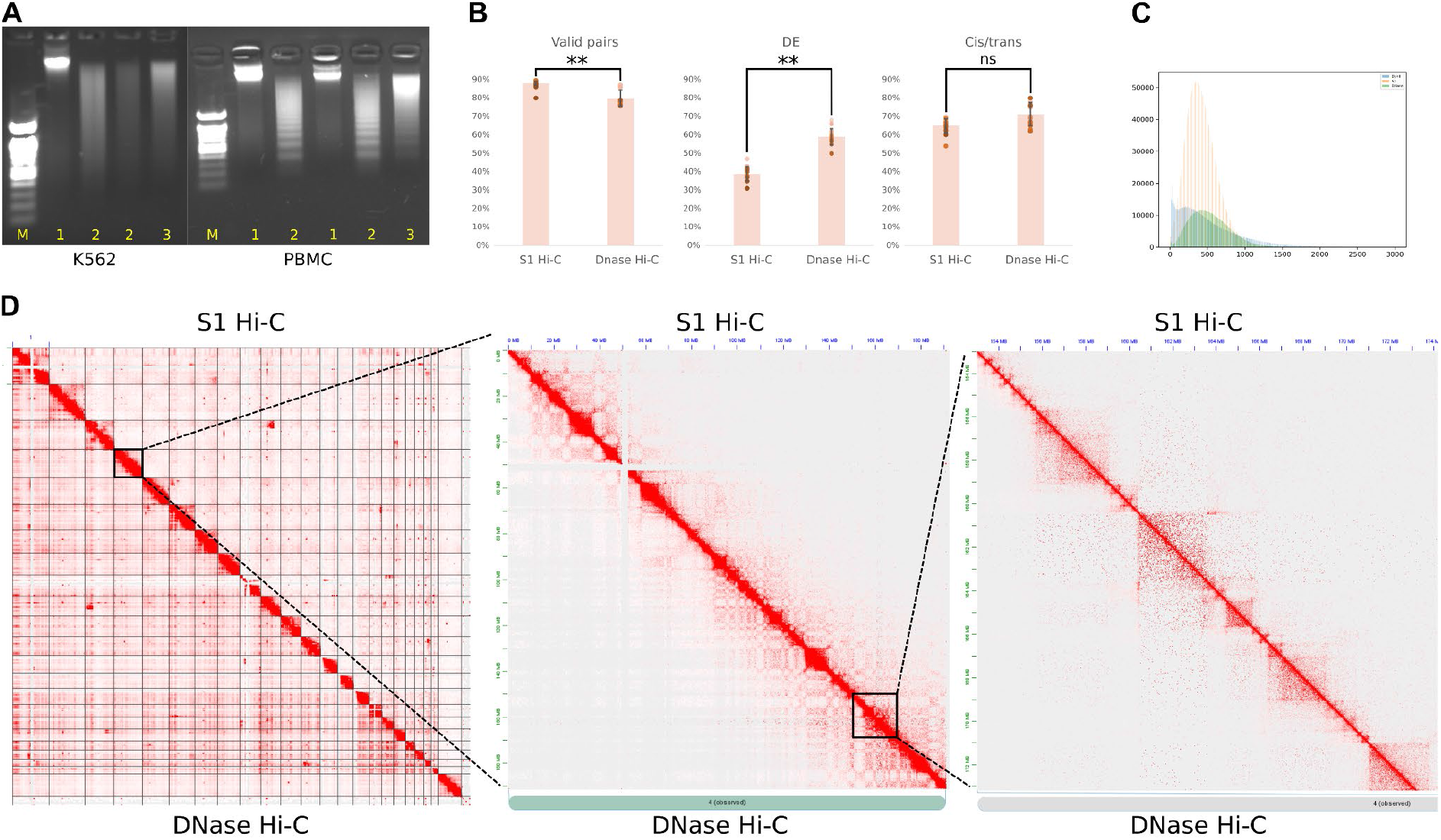
S1 Hi-C protocol allows the generation of high-quality Hi-C maps. (A) Chromatin digestion and ligation of K562 cells and peripheral blood mononuclear cells. Lanes M show a 100 bp DNA ladder. 1 - intact gDNA, 2 - S1 digestion of cross-linked chromatin, 3 - ligation of S1-digested chromatin from lane 2. (B) Quality metrics of S1 Hi-C and DNase Hi-C datasets. Each dot represents an independent Hi-C library preparation; we analyzed 14 DNase Hi-C libraries [11] (protocol with biotin fill-in) and 16 S1 Hi-C libraries. P-values were calculated by using the Mann–Whitney test. (∗∗) indicates p-value < 0.01, (ns) indicates p-value > 0.05. (C) Genome-wide read coverage depth histograms. Blue - DpnII Hi-C; green - S1 Hi-C; yellow - DNAse Hi-C (D) Representative interaction heatmap of K562 cells obtained using DNase Hi-C protocol (below the diagonal line) [11] and S1 Hi-C protocol (above the diagonal line).

Next, we performed shallow sequencing of these samples to access data quality and deeper sequencing for K562 sample to produce Hi-C map (Fig. 1B, D). The quality assessment shows that compared to DNase Hi-C, S1 Hi-C produces similarly high-quality data (Fig. 1B), however the quantity of DEs was lower for S1 Hi-C, resulting in higher overall yield of valid Hi-C pairs. This confirms that the fraction of DE is a consequence of DNAse I nickase activity, and that replacing DNAse I with S1 can reduce the amount of DE fragments.

The DNase I enzyme exhibits a crucial characteristic: its ability to generate Hi-C libraries with a relatively even coverage distribution. Despite a moderate enrichment of A-compartment sequences in DNase I Hi-C libraries [25], this enrichment is less pronounced than that observed at the ends of restriction fragments in conventional Hi-C libraries. To evaluate the potential coverage bias in S1 Hi-C libraries, we computed the distribution of coverage depth across the genome in S1, DNase I, and DpnII Hi-C samples from K562 (Fig. 1C). Our findings revealed that both S1 and DNase I Hi-C maintain relatively even coverage distribution. In contrast, DpnII Hi-C exhibits a bimodal distribution, highlighting the disparity between sequences proximal and distal to restriction sites.

Altogether, our results show that the use of S1 nuclease for chromatin fragmentation makes it possible to achieve the coverage uniformity as in DNase Hi-C and, at the same time, to improve the quality of Hi-C data.

#### Evaluating the Cut Site Distribution in Chromatin Following S1 Nuclease Digestion

Our data suggests that S1 nuclease can be used to digest chromatin, which opens the possibility to apply S1 digestion in various chromatin profiling applications. Although the Hi-C data analysis shows that S1 digestion is fairly uniform across the genome, the complex structure of Hi-C library molecules precludes the precise identification of cut site locations. Therefore, we decided to characterize profiles of S1 nuclease digestion using fixed chromatin, which we fragmented by the enzyme and sequenced. The analysis of the obtained NGS reads revealed that the genomic fragments generated by S1 nuclease start with guanine at their 5’-end approximately two-times more frequent than expected (Fig, 2A). Thus, S1 nuclease has a slight preference to cut the primary strand immediately upstream of the guanine. Interestingly, we did not observe enrichment of cytidine in genomic position immediately before cut site, which would be expected in the case of symmetric cut (Supplemental Fig. 1A, B). This suggests that S1 nuclease probably cleaves DNA strands asymmetrically to form 3’-sticky ends (Supplemental Fig. 1A, B); the length of the overhang can not be determined from our data. These ends could be degraded by S1 nuclease or during subsequent end repair steps of the library preparation.

We next accessed the distribution of DNA fragment lengths obtained by S1 nuclease digestion. Concordant with the electrophoresis gel analysis (Fig. 1A), the NGS data shows that for K562 chromatin there is a minor bias to mono-and di-nucleosomes, whereas for peripheral blood mononuclear cells (PBMC), the specific wavy pattern was more pronounced (Supplemental Fig. 2). The fragment sizes distribution resembled the nucleosomal pattern observed for MNase, thus we reanalyzed data from [26] to compare S1 digestion pattern with the pattern produced by different MNase concentrations (Fig. 2B). MNase digests unprotected linker DNA between nucleosomes, while the DNA protected by the nucleosomes remains intact [27]. Low MNase concentrations generate fragment length distribution corresponding to mono-nucleosome-bound fragments and linker DNA. An increase in MNase concentration leads to a reduction of linker DNA due to its exonuclease activity and fragment length shift to 147 bp (Fig. 2B). Comparison of fragment length distributions suggests that S1 nuclease is more likely to introduce breaks between nucleosomes and has either no or reduced (compared to MNAse) exonuclease activity.

**Figure 2.**
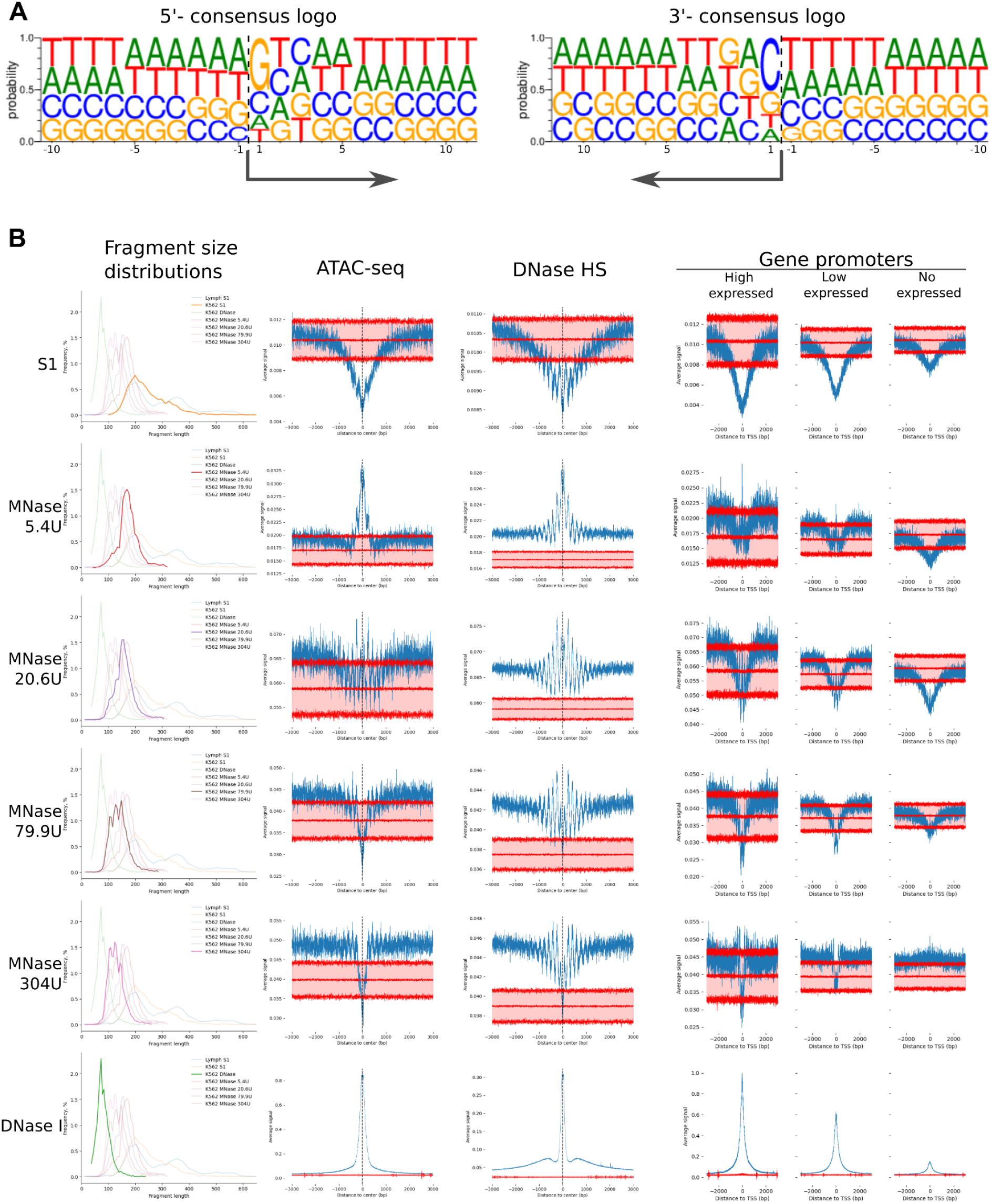
S1 nuclease chromatin digestion pattern. (A) Motif logos represent the sequence specificity of S1 nuclease. Data are shown separately for 5’-(left) and 3’-(right) ends of the digested fragments. In both cases we show the same (reference) strand. The arrow indicates the direction of the sequencing read, and the numbers indicate the distance from the sequenced fragment end: positive numbers for internal (located within the sequenced fragment) nucleotides, negative numbers for external (located outside the sequenced fragment) nucleotides. (B) The fragment size distributions of the mapped paired-end reads and signal distributions at: ATAC-seq peaks, DNase I hypersensitive sites, and TSS for S1 nuclease, different MNAse conditions and DNase in K562 cells.

Next, we aggregated the S1 nuclease, DNAse I, and MNase DNA break location frequencies across annotated open chromatin features: ATAC-seq peaks and DNase I hypersensitive sites (HS) in K562 cells. As DNase I HS and ATAC-seq peaks both align with cis-regulatory elements such as promoters and enhancers of actively transcribed genes, the aggregation of cut sites for these enzymes displays a high degree of concordance (Fig. 2B). The location of MNase cut sites is dependent on enzyme concentration: at low concentrations, the signal heightened across open chromatin regions, implying that these were the first sites accessible to the enzyme. Conversely, at higher MNase concentrations, open chromatin regions were depleted due to elevated digestion of accessible chromatin. The pattern observed for S1 enzyme across DNAse I or ATAC-seq peaks also shows reduced signal in the middle of the peak, followed by gradual increase with nucleosomal pattern (waves). This signal resembles the pattern observed for moderate or high MNAse concentrations; however, nucleosomal pattern was less pronounced than for high MNAse concentration, suggesting that S1 nuclease may not have the strong exonuclease activity required to digest linker DNA.

Nucleosome positioning and chromatin accessibility shows non-random pattern across gene promoters, with the level of accessibility correlating with the gene expression. We analyzed the chromatin cut frequency across transcription start sites (TSS) stratifying genes by the expression level (Fig. 2B). Both S1 nuclease and MNAse show decreased fragment ends frequency in the open chromatin region, presumably because these regions are over-digested, producing very small fragments (or even digested into individual nucleotides), that can not be captured by sequencing. The reduced signal region was broader than for MNAse data, and does not show a clear nucleosome pattern.

Drawing from these findings, we posit that S1 nuclease exhibits greater activity on DNA bound to nucleosomes than MNase, while its exonuclease activity to digest linker DNA is either lower or non-existent. This results in a more uniform distribution of S1 nuclease cut sites in comparison to MNase. In relation to DNAse, S1 nuclease presents a decreased representation of fragment ends within open chromatin regions, a pattern that may be attributable to the presence of exonuclease activity, or possibly due to high endonuclease activity that reduces these loci into fragments too minuscule for detection. Finally, S1 nuclease shows slight preference towards cleavage of guanine 5-phosphate bonds, leaving a 3’-overhang on the complementary strand. Despite these preferences, the overall pattern of S1 cuts is relatively uniform (compared to DNAse I or MNAs digestion) and thus allows studying both open and closed chromatin.

## Discussion

Developing new enzymes for chromatin profiling is essential for genomics. Here, we show that S1 nuclease can be used to cut fixed chromatin. We note that we did not profile all digestion conditions thoroughly: varying enzyme concentration, incubation times, digestion buffers, temperature, and other conditions may result in different digestion patterns. This more detailed profiling is an important direction of further research, because for DNAse I and MNAse it has been shown that varying reaction conditions significantly affect the specificity and activity of these enzymes [28].

One important application of the current study is using S1-nuclease for Hi-C analysis. Sequence-agnostic digestion with S1 nuclease is important for genome assembly or structural variant detection, where the resolution of breakpoints is limited by a frequency of genomic location of digestion sites. We developed a robust and efficient protocol for S1 Hi-C analysis of PBMC, which can be used to capture structural variants in cells and tissues in humans. Future applications can extend S1 Hi-C protocols for genome assembly and haplotyping.

## Data access

Sequencing data generated in this study are accessible via the NCBI BioProject PRJNA983247

## Competing interests statement

Authors declare no competing interest.

## Acknowledgment

We acknowledge the Ministry of Science and Higher Education of the Russian Federation (state project FWNR-2022-0019) for providing access to the computational facilities. The access to public resources and datasets was provided by the Novosibirsk State University, supported by the Ministry of Education and Science of Russian Federation, grant #2019-0546 (FSUS-2020-0040). The original manuscript text was composed by authors; proofreading was done with assistance of the chatGPT version 4. The text was corrected and edited by authors after chatGPT proofreading.

## Funding

S1 Hi-C library preparation, sequencing and analysis was supported by RSCF Grant no. 22-24-00190.

## Author contributions

VF designed and supervised the study; MG performed all the wet-lab experiments; PA, SA, TN performed the computational data analysis; BU, KA, KE, RO, FM performed NGS sequencing; VF, MG wrote the manuscript; all the authors revised the manuscript. All authors read and approved the final manuscript.

## Ethics approval and consent to participate

This study was performed in line with the principles of the Declaration of Helsinki. Approval was granted by the local Research Ethics Committee of the Research Institute of Medical Genetics, Tomsk National Research Medical Center (Date 27.07.2017/No 106). All the study participants provided informed consent.

## Notes

### Competing Interest Statement

The authors have declared no competing interest.

## References

1. Klein, D.C.; Hainer, S.J. Genomic Methods in Profiling DNA Accessibility and Factor Localization. Chromosome Res 2020, 28, 69–85, doi:10.1007/s10577-019-09619-9.

2. Buenrostro, J.D.; Giresi, P.G.; Zaba, L.C.; Chang, H.Y.; Greenleaf, W.J. Transposition of Native Chromatin for Fast and Sensitive Epigenomic Profiling of Open Chromatin, DNA-Binding Proteins and Nucleosome Position. Nat Methods 2013, 10, 1213–1218, doi:10.1038/nmeth.2688.

3. Fishman, V.; Battulin, N.; Nuriddinov, M.; Maslova, A.; Zlotina, A.; Strunov, A.; Chervyakova, D.; Korablev, A.; Serov, O.; Krasikova, A. 3D Organization of Chicken Genome Demonstrates Evolutionary Conservation of Topologically Associated Domains and Highlights Unique Architecture of Erythrocytes’ Chromatin. Nucleic Acids Research 2019, 47, 648–665, doi:10.1093/nar/gky1103.

4. Gridina, M.; Taskina, A.; Lagunov, T.; Nurislamov, A.; Kulikova, T.; Krasikova, A.; Fishman, V. Comparison and Critical Assessment of Single-Cell Hi-C Protocols. Heliyon 2022, 8, e11023, doi:10.1016/j.heliyon.2022.e11023.

5. Fishman, V.S.; Salnikov, P.A.; Battulin, N.R. Interpreting Chromosomal Rearrangements in the Context of 3-Dimentional Genome Organization: A Practical Guide for Medical Genetics. Biochemistry (Mosc) 2018, 83, 393–401, doi:10.1134/S0006297918040107.

6. Kabirova, E.; Nurislamov, A.; Shadskiy, A.; Smirnov, A.; Popov, A.; Salnikov, P.; Battulin, N.; Fishman, V. Function and Evolution of the Loop Extrusion Machinery in Animals. Int J Mol Sci 2023, 24, 5017, doi:10.3390/ijms24055017.

7. Gridina, M.; Fishman, V. Multilevel View on Chromatin Architecture Alterations in Cancer. Front Genet 2022, 13, 1059617, doi:10.3389/fgene.2022.1059617.

8. Lukyanchikova, V.; Nuriddinov, M.; Belokopytova, P.; Taskina, A.; Liang, J.; Reijnders, M.J.M.F.; Ruzzante, L.; Feron, R.; Waterhouse, R.M.; Wu, Y.; et al. Anopheles Mosquitoes Reveal New Principles of 3D Genome Organization in Insects. Nat Commun 2022, 13, 1960, doi:10.1038/s41467-022-29599-5.

9. Maslova, A.; Plotnikov, V.; Nuriddinov, M.; Gridina, M.; Fishman, V.; Krasikova, A. Hi-C Analysis of Genomic Contacts Revealed Karyotype Abnormalities in Chicken HD3 Cell Line. BMC Genomics 2023, 24, 66, doi:10.1186/s12864-023-09158-y.

10. Mozheiko, E.A.; Fishman, V.S. Detection of Point Mutations and Chromosomal Translocations Based on Massive Parallel Sequencing of Enriched 3C Libraries. Russian Journal of Genetics 2019, 55, doi:10.1134/S1022795419100089.

11. Gridina, M.; Mozheiko, E.; Valeev, E.; Nazarenko, L.P.; Lopatkina, M.E.; Markova, Z.G.; Yablonskaya, M.I.; Voinova, V.Y.; Shilova, N.V.; Lebedev, I.N.; et al. A Cookbook for DNase Hi-C. Epigenetics Chromatin 2021, 14, 15, doi:10.1186/s13072-021-00389-5.

12. Belton, J.-M.; McCord, R.P.; Gibcus, J.; Naumova, N.; Zhan, Y.; Dekker, J. Hi-C: A Comprehensive Technique to Capture the Conformation of Genomes. Methods 2012, 58, 10.1016/j.ymeth.2012.05.001, doi:10.1016/j.ymeth.2012.05.001.

13. Rao, S.S.P.; Huntley, M.H.; Durand, N.C.; Stamenova, E.K.; Bochkov, I.D.; Robinson, J.T.; Sanborn, A.L.; Machol, I.; Omer, A.D.; Lander, E.S.; et al. A 3D Map of the Human Genome at Kilobase Resolution Reveals Principles of Chromatin Looping. Cell 2014, 159, 1665–1680, doi:10.1016/j.cell.2014.11.021.

14. Lafontaine, D.L.; Yang, L.; Dekker, J.; Gibcus, J.H. Hi-C 3.0: Improved Protocol for Genome-Wide Chromosome Conformation Capture. Curr Protoc 2021, 1, e198, doi:10.1002/cpz1.198.

15. Ramani, V.; Cusanovich, D.A.; Hause, R.J.; Ma, W.; Qiu, R.; Deng, X.; Blau, C.A.; Disteche, C.M.; Noble, W.S.; Shendure, J.; et al. Mapping 3D Genome Architecture through in Situ DNase Hi-C. Nat Protoc 2016, 11, 2104–2121, doi:10.1038/nprot.2016.126.

16. Hsieh, T.-H.S.; Weiner, A.; Lajoie, B.; Dekker, J.; Friedman, N.; Rando, O.J. Mapping Nucleosome Resolution Chromosome Folding in Yeast by Micro-C. Cell 2015, 162, 108–119, doi:10.1016/j.cell.2015.05.048.

17. Akgol Oksuz, B.; Yang, L.; Abraham, S.; Venev, S.V.; Krietenstein, N.; Parsi, K.M.; Ozadam, H.; Oomen, M.E.; Nand, A.; Mao, H.; et al. Systematic Evaluation of Chromosome Conformation Capture Assays. Nat Methods 2021, 18, 1046–1055, doi:10.1038/s41592-021-01248-7.

18. Shishido, K.; Ando, T. Site-Specific Fragmentation of Bacteriophage T5 DNA by Single-Strand-Specific S1 Endonuclease. Biochim Biophys Acta 1975, 390, 125–132, doi:10.1016/0005-2787(75)90015-5.

19. Kouzine, F.; Wojtowicz, D.; Baranello, L.; Yamane, A.; Nelson, S.; Resch, W.; Kieffer-Kwon, K.-R.; Benham, C.; Casellas, R.; Przytycka, T.M.; et al. Permanganate/S1 Nuclease Footprinting Reveals Non-B DNA Structures with Regulatory Potential across a Mammalian Genome. Cell Syst 2017, 4, 344–356.e7, doi:10.1016/j.cels.2017.01.013.

20. Mimitou, E.P.; Yamada, S.; Keeney, S. A Global View of Meiotic Double-Strand Break End Resection. Science 2017, 355, 40–45, doi:10.1126/science.aak9704.

21. Matos-Rodrigues, G.; Wietmarschen, N. van; Wu, W.; Tripathi, V.; Koussa, N.C.; Pavani, R.; Nathan, W.J.; Callen, E.; Belinky, F.; Mohammed, A.; et al. S1-END-Seq Reveals DNA Secondary Structures in Human Cells. Molecular Cell 2022, 82, 3538–3552.e5, doi:10.1016/j.molcel.2022.08.007.

22. Ramírez, F.; Ryan, D.P.; Grüning, B.; Bhardwaj, V.; Kilpert, F.; Richter, A.S.; Heyne, S.; Dündar, F.; Manke, T. DeepTools2: A next Generation Web Server for Deep-Sequencing Data Analysis. Nucleic Acids Research 2016, 44, W160–W165, doi:10.1093/nar/gkw257.

23. Harris, C.R.; Millman, K.J.; van der Walt, S.J.; Gommers, R.; Virtanen, P.; Cournapeau, D.; Wieser, E.; Taylor, J.; Berg, S.; Smith, N.J.; et al. Array Programming with NumPy. Nature 2020, 585, 357–362, doi:10.1038/s41586-020-2649-2.

24. Hunter, J.D. Matplotlib: A 2D Graphics Environment. Computing in Science & Engineering 2007, 9, 90–95, doi:10.1109/MCSE.2007.55.

25. Chandradoss, K.R.; Guthikonda, P.K.; Kethavath, S.; Dass, M.; Singh, H.; Nayak, R.; Kurukuti, S.; Sandhu, K.S. Biased Visibility in Hi-C Datasets Marks Dynamically Regulated Condensed and Decondensed Chromatin States Genome-Wide. BMC Genomics 2020, 21, 175, doi:10.1186/s12864-020-6580-6.

26. Mieczkowski, J.; Cook, A.; Bowman, S.K.; Mueller, B.; Alver, B.H.; Kundu, S.; Deaton, A.M.; Urban, J.A.; Larschan, E.; Park, P.J.; et al. MNase Titration Reveals Differences between Nucleosome Occupancy and Chromatin Accessibility. Nat Commun 2016, 7, 11485, doi:10.1038/ncomms11485.

27. Noll, M.; Thomas, J.O.; Kornberg, R.D. Preparation of Native Chromatin and Damage Caused by Shearing. Science 1975, 187, 1203–1206, doi:10.1126/science.187.4182.1203.

28. Melgar, E.; Goldthwait, D.A. Deoxyribonucleic Acid Nucleases. II. The Effects of Metals on the Mechanism of Action of Deoxyribonuclease I. J Biol Chem 1968, 243, 4409–4416.

